# Physics-guided design of intrinsically disordered proteins

**DOI:** 10.64898/2026.05.29.728696

**Authors:** Neha Tyagi, Jackson Boodry, Vita Chou, Wilton T. Snead, Krishna Shrinivas

## Abstract

Intrinsically disordered protein regions (IDPs) are found across the tree of life and characterized by the lack of a stable 3D fold, encoding function through a vast ensemble of conformations. This plasticity makes rational design of IDPs challenging. Physics-based approaches capturing distinct aspects of sequence composition, charge patterning, and molecular interactions have emerged as powerful predictors of ensemble-derived properties. Here, we present a machine learning framework for proteome-scale de novo IDP design by rationally inverting physics-based models. We first program IDPs to tunably sense and respond to diverse biophysical cues and show that IDP ensembles can directly encode complex signal processing, including threshold detection, bandpass filtering, and Boolean-type multi-input logic. We next engineer multicomponent IDP mixtures with tailored emergent condensate properties, including layering and number of phases, compositional specificity, and RNA-dependent remodeling of structure and composition. Finally, we demonstrate designed IDPs that selectively partition into or deplete from biological condensates in living cells. Together, our framework establishes a flexible and scalable strategy for design of ensemble-derived and collective properties in dynamic biomolecules.

## Introduction

Intrinsically disordered proteins and protein regions (collectively IDPs) are widespread throughout the tree of life and play central roles in many key cellular processes^1–3^. Unlike folded proteins that have largely static three dimensional structures, IDPs occupy a heterogeneous ensemble of interconverting conformations^4,5^ that shapes their functions. Since the structural ensemble of IDPs is highly dependent on physicochemical and biological contexts, many IDPs function as cellular sensors ^6,7^. IDPs are capable of dynamic, often multivalent, interactions, and thus play key roles in integrating information across multiple binding partners as well as shaping subcellular localization, particularly into biomolecular condensates. Mutants of IDPs that disrupt or alter ensemble-derived properties are increasingly understood to correlate with pathology^8^. Given their prevalence and functional diversity, there is growing interest to design IDPs with specified ensemble properties.

The rational design of folded proteins has dramatically accelerated in recent years, built upon inverting increasingly accurate predictors of sequence-structure maps such as AlphaFold and ESMFold^9,10^. By contrast, IDP ensembles are inherently more complex, are naturally sensitive to sequence, environment, and biological context, and not described by a single native structure^7^. Physics-based simulations and theoretical models are emerging as powerful tools to predict IDP ensemble properties. However, there are many models that each emphasize distinct sequence features, can be expensive to compute, and remain challenging to invert^11^. Thus, there is interest in developing algorithms that can scalably and flexibly design IDPs with a diverse range of biologically-motivated, context-dependent behavior.

Here we develop a physics-centered machine learning approach to design intrinsically disordered proteins with tailored ensemble properties by inverting physics-based models. Our framework efficiently optimizes IDP sequences and is scalable, enabling proteome-level design within a minute on GPUs. We demonstrate the generation of de novo IDPs with a number of complex phenotypes: first, we design IDPs that are stimuli-responsive, exploiting the sensitivity of IDP ensembles to their environment to enable nonlinear environmental sensing and multi-stimuli information processing. Second, we program multicomponent IDP mixtures to condense and drive emergent behaviors, including coexistence of multiple phases, layered architectures, and RNA-dependent remodeling of composition. Third, we design IDPs that selectively localize or exclude from specific biological condensates. We then test and validate our predictions through localization assays in living cells. Overall, our work outlines a scalable, physics-based platform for IDP design with diverse ensemble-derived functional properties.

## Results

### Model framework

Physics-derived models have recently been developed to predict thermodynamic properties of IDPs directly from sequence and physical parameters^12–14^(Fig. 1A). These models naturally accommodate changing biophysical stimuli such as salt concentration and phosphorylation status and are parameterized by force fields^15–20^ coarse-grained at the 1AA=1bead resolution.

**Figure 1.**
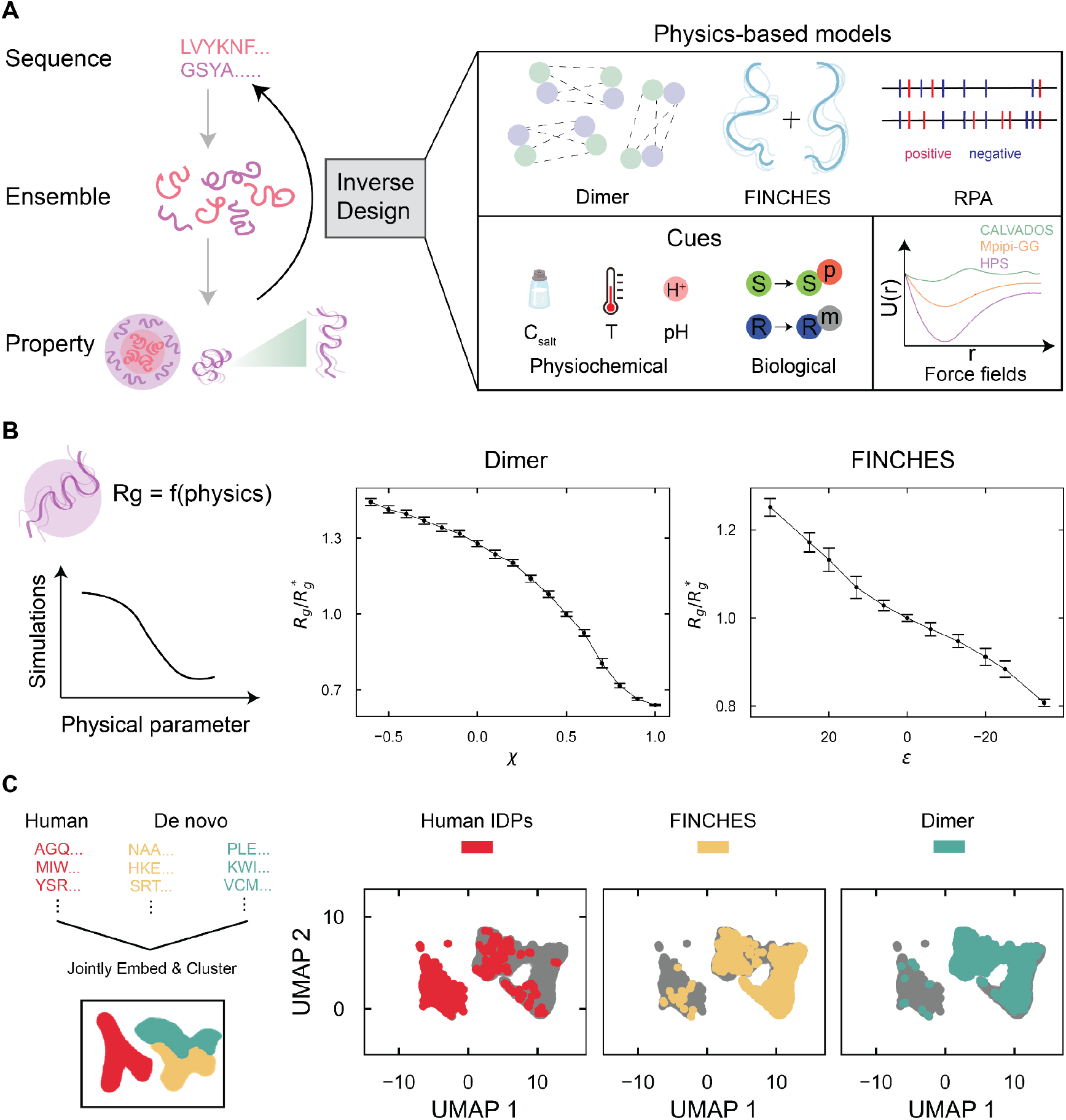
Framework for physics-based design of IDPs. **A**. The computational framework for IDP design inverts sequence-ensemble-property relationships (left) predicted by physics-based approaches (right) and flexibly incorporates distinct models, cues, and choice of forcefields (right). **B**. The left panel depicts a schematic showing trends in simulation predictions of Rg with physics-model derived parameters. Simulation-predicted Rg values (y axis) versus self-interaction values (x-axis) for dimer (center) and FINCHES (right) models. The Rg values are normalized to the average Rg* at χ=0.5 and ε=0. Each dot is the average over simulations of 25 independent designs, error bars are standard deviations, and the black line is a guide. **C**. Schematic description (left) of sequence analysis procedure. (right) UMAP visualization on joint sequence embedding, with points colored according to their source. Human IDPs are in red, FINCHES-designed sequences in yellow, and dimer model-designed sequences in green. In each plot, grey points correspond to the rest of the dataset.

They differ in how they relate sequence features to ensemble-averaged properties (Fig. 1A) as described briefly (see SI Note 2 for an expanded discussion). The dimer model describes each sequence (GSGY for example) by the number and type of amino acid doublets (1GS, 1SG, 1GY) to describe local sequence connectivity. Treating a sequence as a bag of dimers, interactions between pairs of IDPs are generated from this encoding through physics-derived weighting rules. The FINCHES model follows a similar approach but differs in that IDPs are described by their amino acid composition (2G, 1S, 1Y) i.e., as bags of monomers, and empirically determined local sequence context rules are additionally incorporated. Finally, the random phase approximation (RPA) model combines a monomer-style treatment with explicit consideration of sequence-wide context for all charged residues. Predictions from these models have been shown to correlate with experimental or simulation data, but not necessarily for the same set of sequences and depend on the force field of choice^12–14^. Our goal is to develop a general computational framework to design IDPs by flexibly inverting these models.

To achieve this, we first implemented a differentiable physics-based framework that is amenable to gradient-based design^11,21,22^. Across models (dimer, FINCHES, RPA), force fields (CALVADOS, Mpipi-GG, HPS), and diverse sequences, we confirmed that predictions matched their original formulations (Fig. S1). Second, we employ a probabilistic sequence representation that is amenable to gradient-based optimization and progressively annealed to produce discrete amino acid sequences^23–25^ (see SI Note 1). Third, we enforce that final designed sequences are intrinsically disordered (>80% disordered by Metapredict^26^ V3). Finally, a loss function that encodes the desired property is specified to design *de novo* IDP sequences using standard stochastic gradient-descent optimization techniques^27^ (see SI Note 1).

Together, these enable our design framework to perform de novo IDP generation across distinct model priors and flexible to design choice. Our framework is quick (few seconds/sequence), works on CPUs and GPUs, and scales to proteome-scale generation of IDPs (Fig. S2, 10000 sequences in <1 minute). Designs were validated through explicit molecular dynamics (MD) simulations that directly sample the IDP ensemble (Figs5. 1-3) and also through experiments in cells for a subset of designs (Fig. 4).

### Flexible design of IDP ensemble dimensions

The ensemble-averaged physical dimensions of an IDP, particularly the radius of gyration (Rg), correlates with binding or phase behavior^28,29^ and thus our first design objective. While physics-based models do not directly predict Rg, model-derived self-interaction parameters (χ_ii_ for the dimer model, ε for FINCHES, χ_RPA_ for the RPA model) should correlate with Rg (Fig. 1B). To evaluate this, we used the dimer model parameterized by the CALVADOS force field to generate IDP sequences across a range of self-interaction parameter (χ) values and calculated their corresponding Rg from MD simulations^30^ (see SI Notes 1 and 2). We find a strong concordance between physics-derived χ and simulation-derived Rg (Fig. 1B, centre) for our designed sequences to <5%. We identify similar trends for FINCHES (Fig. 1B, right) and RPA models (Fig. S3) as well as for other force fields (Fig. S4). While these results indicate that physics-derived models are good approximations of explicit simulations, it is important to note that these may not be exact everywhere in sequence-space. A key advantage of designing over physics-based models is their scalability and speed (1000s of sequences/s) in contrast to directly inverting molecular simulations^11^ (1 seq/ 10,000s) (Fig. S2). Together, our approach enables rapid, rational design of IDPs with desired ensemble-averaged Rgs.

We next sought to assess the sequence diversity of de novo IDPs and contrast with the human IDPome. Towards this, we first constructed a dataset containing all human IDPs of lengths between 30 and 70 AA (N∼8,500). We then constructed de novo IDPomes (N∼8,500 proteins, L=50AA/IDP) using two physics models (dimer, FINCHES) with distribution of self-interactions (χ, ε) matched to human IDPome (see SI Note 3). This allows us to dissect sequence diversity while controlling for length, number, and interaction statistics of the human IDPome and only requires a minute on a standard GPU for IDP library generation (Fig. S2). For each library, we generated sequence representations using the ESM2 protein language model^31,32^ and assessed diversity through unsupervised K-Means clustering (Figs. S5 and S6) and visualized through 2D UMAP embeddings (Fig. 1C) (See SI Note 3). Our analysis shows that designed sequences are largely distinct in comparison to the human proteome for both physics models (Fig. 1C, Fig. S5). When repeating a similar analysis without matching for interaction statistics of human IDPs, sequences from different models largely cluster separately (Fig. S6). We find similar results when we represent sequences through the dimer frequency content (Figs. S5, S6). Together, these suggest that designing over physics-based models largely generates IDPs that are distinct from human IDPs. For the rest of the manuscript, unless otherwise specified, we will use the dimer model with the CALVADOS force field for de novo IDP design.

### Multiplexed and nonlinear signal processing by stimuli-responsive designed IDPs

Many naturally occurring IDPs sense and respond to physicochemical and biological stimuli^7,33,34^ such as cellular salt concentrations, pH levels, and phosphorylation state by changing global- or local-chain conformations. Motivated by this, we next sought to design IDP sensors. We define sensor function as large changes in Rg in response to changes in environmental stimuli or post-translational modifications (see SI Note 1). We first generated IDPs that contract in response to a defined increase in salt concentration from 150 to 450 mM. Our designed IDPs exhibited large changes in Rg (∼25%) (Fig. 2A), comparable to previously reported salt sensors^11^ that required 1000-10000x more time to design. By solely changing the loss, we generated a panel of contractor and expander sensors of diverse biophysical stimuli (Table S1) albeit with cue-dependent dynamic range in chain conformations.

**Figure 2.**
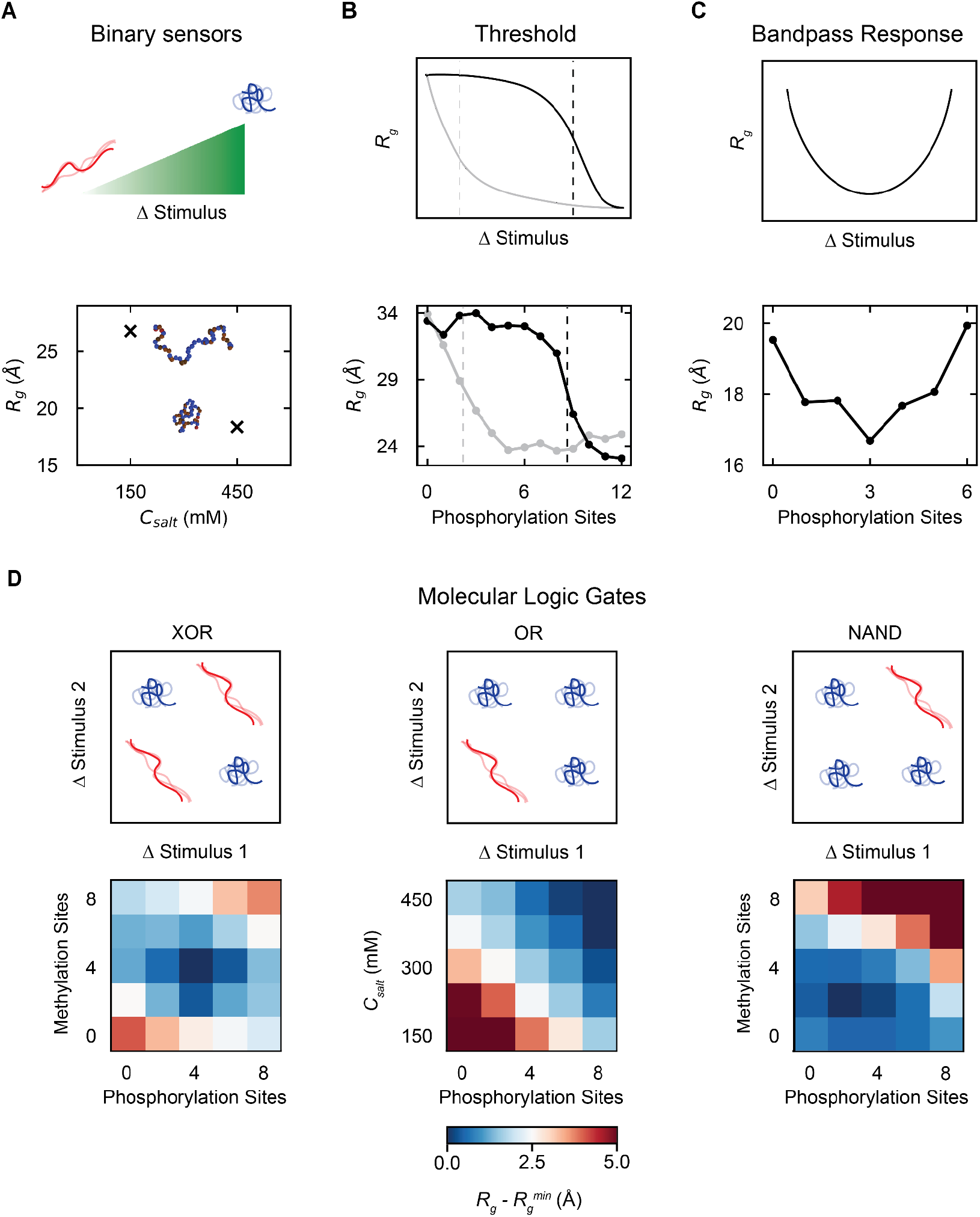
Multiplexed and nonlinear signal processing by stimuli-responsive IDPs. **A**. Top panel depicts the objective of contraction upon stimuli. The bottom panel shows simulated Rg of a designed IDP (y-axis) in response to changing salt concentration (x-axis), and the insets highlight characteristic conformations. **B**. Top panel schematically depicts the objective of threshold sensing IDPs. Bottom panel shows simulation-derived Rg (y-axis) in response to increasing phosphorylation levels (x-axis) for two designs that exhibit a threshold response at low (grey) or high (black) phosphorylation. **C**. Top panel schematically depicts the objective of bandpass sensors. Bottom panel shows simulation-derived Rg (y-axis) versus phosphorylation level (x-axis). **D**. Engineered IDP sensors that approximate boolean-type computations in their ensemble dimensions in response to two distinct stimuli with objectives schematized above and simulation derived Rg below. The specific computations performed by each IDP are XOR (left), OR (center), and NAND (right).

Sensing, transduction, and information processing are typically enacted by biochemical signaling cascades with multiple enzymes^35,36^ i.e. modular cascades with different molecules that perform dedicated roles. Biomolecules can also directly sense (by changing conformational ensemble) and respond (altered ensemble-derived property like binding or condensation), a form of physical signal processing exemplified by allostery^37,38^ in folded domains as well as IDP sensors^1^. For example, the prion-like IDP domain of the ELF3 protein functions as a thermosensor in plants (*Arabidopsis Thalliana)*, and is characterized by a sharp transition in organismal phenotypes above and below a critical temperature^39^. Motivated by this, we next asked whether such physical computations can be directly engineered into the sequence of an IDP by sculpting the expansion-contraction response. We first generated threshold sensors, i.e. IDPs that undergo a rapid change in Rg only beyond a threshold value of stimulus. This is highlighted in Fig. 2B by IDPs that effectively “count” phosphorylation status into an ensemble-derived Rg, with each sequence displaying a distinct threshold. Motivated by biological signalling pathways that perform complex information processing tasks^36,40,41^, we next sought to design non-linear IDP responses. We designed a bandpass sensing IDP that adopts lower Rg values at intermediate phosphorylation levels but higher Rg otherwise (Fig. 2C), thus effectively “filtering” out low or high phosphorylation levels. Collectively, these examples highlight that tuning stimuli-responsive behavior is a promising avenue for IDP-mediated physical signal processing.

A key feature of biological signaling is the ability to respond and integrate multiple environmental stimuli. A prominent example is Pab1, a protein whose unfolding and condensation enables sensing of both temperature and pH in yeast^42^. We sought to encode this kind of multidimensional information processing into IDPs. To achieve this requires constraints on the ensemble-statistics across a range of conditions, a challenging multi-objective design problem that can be tackled by our framework. We first model salt concentration and extent of phosphorylation simultaneously. We generated an IDP that implements an OR-gate-like logic: the sequence adopts an expanded conformation when unphosphorylated and at physiological salt concentration, and contracts under all other combinations of conditions (Fig. 2D, center). We additionally designed IDPs that implement both XOR- and NAND-like logic in response to two environmental stimuli inputs (Fig. 2D, left and right) and highlight how these designs can generalize to more stimuli and computations (Fig S7). Together, these results show that single IDP sequences can be designed to encode and perform boolean-type molecular computations that directly integrate multiple cues.

### Programming condensate composition, architecture, and RNA-dependent reorganization in multicomponent IDP mixtures

IDPs, often with other biomolecules, self-organize by phase transitions into biomolecular assemblies called condensates. Condensates with IDPs are ubiquitous in living cells^43^ and are increasingly harnessed for synthetic, programmable biomaterial design^44,45^. Unlike oil-water mixtures or protein aggregates, condensates exhibit diverse and functional organization, including concentrating multiple species^46^, forming coexisting phases with differing compositions, selectively recruiting or excluding molecules, and exhibiting spatially layered architectures that are thought to aid in organizing biochemical pathways. Because these emergent properties arise from a network of molecular interactions^47^, we next asked whether we could design multicomponent mixtures of de novo IDPs that self-organize into condensates with programmable properties. We then extend this design to account for the presence of RNA, a common feature of biomolecular condensates whose presence has been shown to influence condensate architecture in vivo^48,49^. Towards this, we extended our framework to allow for joint optimization of multiple IDP sequences. We then seek to tailor the *entire* network of molecular interactions, comprising both self (χ_ii_ for IDP *i*) and cross interactions (χ_ij_ for IDPs *i* and *j*) to achieve a desired condensate property (see SI Note 1). Once designed, we evaluate the resulting phase behavior using direct coexistence (slab) molecular dynamics simulations, focusing on equimolar mixtures of IDPs unless otherwise noted.

We demonstrate the flexibility of our framework by designing multicomponent mixtures (for e.g. requiring joint optimization of 4 sequences with 10 interaction parameters) to self-organize into condensates with diverse properties. First, we design a mixture to show multiphase coexistence, forming two compositionally-unique condensates that do not mix (Fig. 3B). Second, we program spatially heterogeneous condensates, where a set of three de novo IDPs condense into 3 nesting layers (Fig. 3C), qualitatively analogous to the 3-layered architecture of nucleoli^50^. Third, we encode condensate specificity, where a mixture of IDPs form a condensate that specifically only enriches 2 out of 4 species (Fig. 3D) and excludes the rest. Finally, we demonstrate the design of two de novo IDPs whose phase behavior is dependent on the presence of RNA (Fig. 3E). Using polyU-50 as our example RNA molecule, in the absence of RNA the two designed IDPs form a condensed, mixed droplet (Fig. 3E, center). Introduction of RNA promotes the formation of two phases within the larger condensate: one RNA-rich phase enriched with one designed IDP and an RNA-depleted phase enriched with the other designed IDP (Fig. 3E, right). We further show that our framework accommodates varying numbers of IDPs and condensate properties (Fig. S8), although we note that the optimization constraints scale quadratically with the number of species, limiting generalization to arbitrary numbers. Together, our results show that we can optimize and rationally design multiple interacting IDPs to achieve programmable emergent condensate properties.

**Figure 3.**
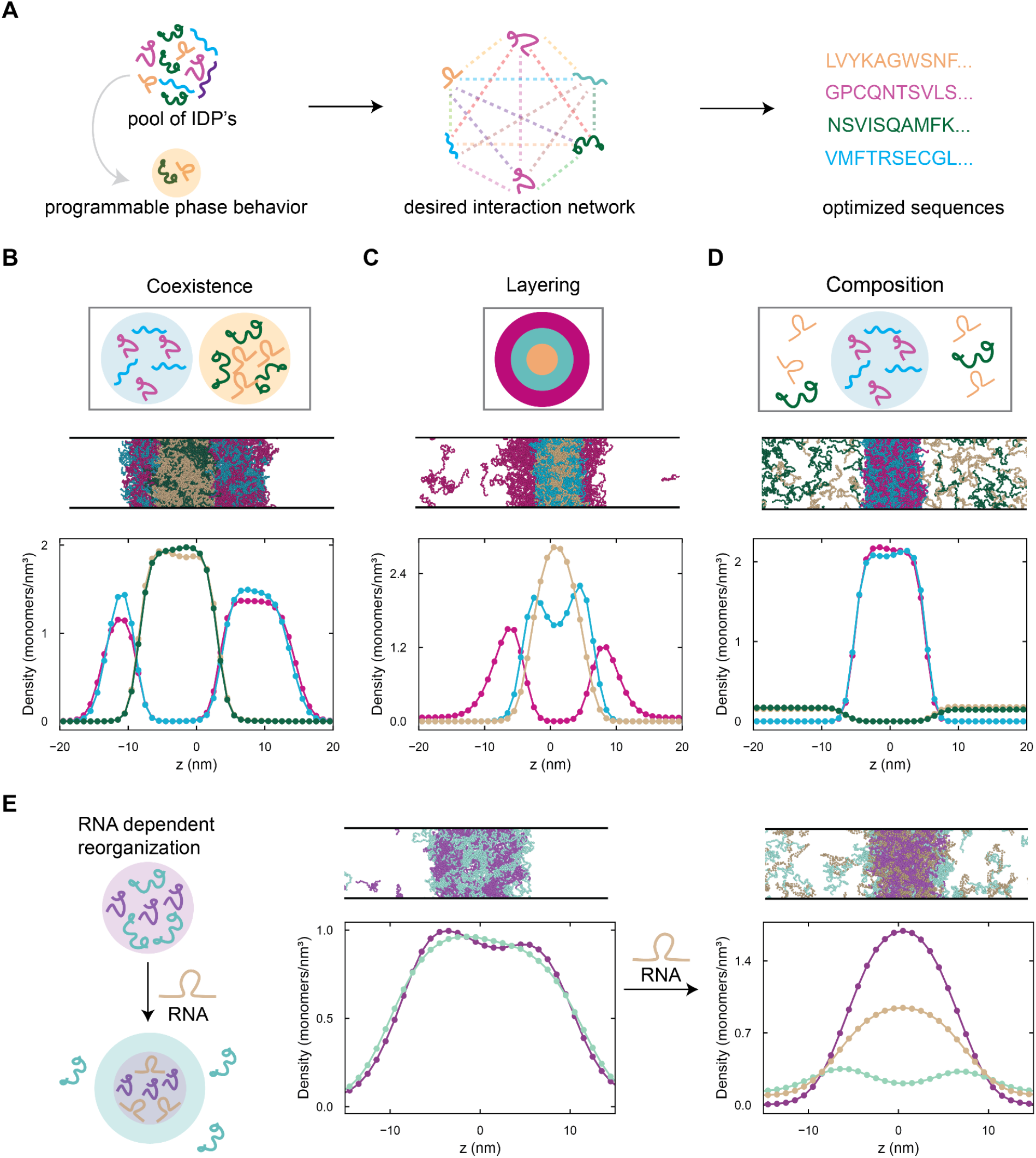
Programming condensate composition, architecture, and RNA-dependent organization in multicomponent IDP mixtures. **A**. Schematic overview of programmable multicomponent, multiphase condensate design through tuning IDP interaction network. **B-D**., Schematic of desired phase behavior (top), snapshot from slab molecular dynamics simulation (middle), and quantified density profiles from simulations (bottom) is shown. The condensate designs are multiphase coexistence (B), condensate layering (C), and compositional specificity (D). **E**. Schematic of RNA-dependent condensate reorganization (left). The two designed IDPs mix in a single condensed phase (center) that remodels into an RNA-rich condensate with different composition upon presence of RNA (right).

**Figure 4.**
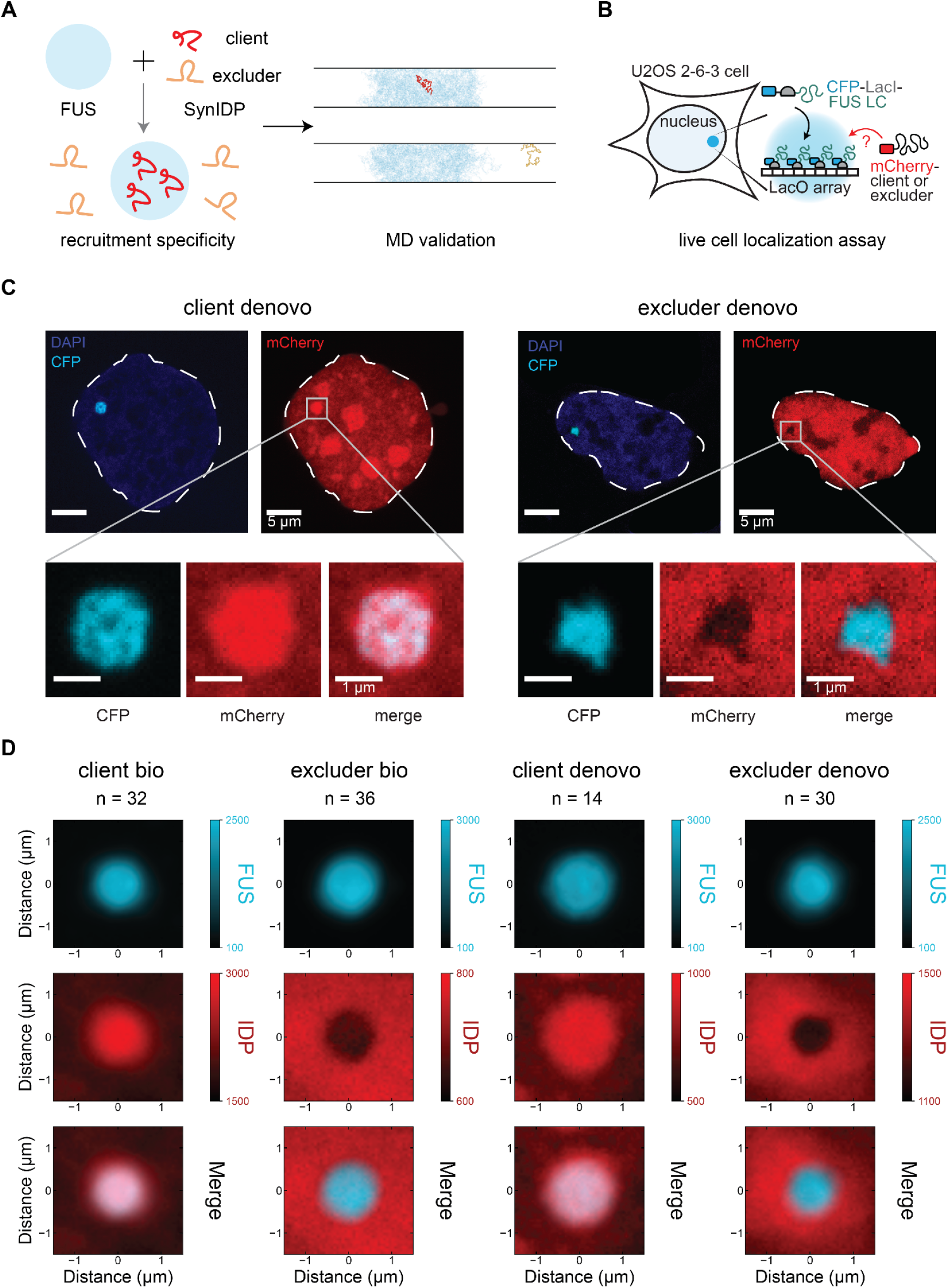
Engineering IDP localization to cellular compartments. **A**. The design objective (left) for IDPs that selectively enrich (client) or deplete (excluder) in FUS LC condensates. Representative snapshots (right) of FUS LC clients and excluders from molecular dynamics simulation for computational validation. **B**. Schematic of live-cell assay with CFP- and LacI-tagged FUS LC (blue) co-expressed with mCherry-tagged clients or excluders (red) in U2OS cells. **C**. Representative composite images of a cell with conditions for designed clients (left) and excluders (right). Each composite has a nuclear outline (dashed lines), DAPI (blue), CFP (FUS LC), and mCherry (IDP) channels, as well as a zoom-in and merge focused on the condensate. **D**. Averaged images for de novo and biological (bio) clients and excluders centered on a 3 × 3 µ*m* box around each FUS LC condensate, averaged over the indicated number of images. First row is the CFP channel (FUS LC), second row is mCherry (IDP), and third row is merge.

### Engineering IDP localization to cellular compartments

Multivalent interactions mediated by biological IDPs contribute to selective recruitment of molecules into condensates that, in turn, shape downstream functions^51^. For example, the disordered region of MED1 partitions positive but not negative regulators of gene expression into transcriptional condensates^52^. Motivated by this, we next sought to design IDPs that selectively enrich (client) or deplete (excluder) from a condensate rich in a biologically relevant scaffold protein. For the scaffold, we focused on the low-complexity domain of the protein FUsed in Sarcoma (FUS LC here on) that has been extensively characterized in the literature^17,53,54^, and has been shown to modulate IDP recruitment in vitro^55^. To design clients (or excluders), we optimized the cross-interaction parameter between a designed IDP and the FUS LC (χ_i,FUS_) to favor (or disfavor) heterotypic interactions (see SI Note 1, Fig. 4A). We validated our designs through explicit slab simulations, finding that χ_i,FUS_ has a direct correlation with the extent of protein localization (Fig. S9). To ensure that desired behavior arose from sequence-intrinsic properties, we used the same validation criteria to identify length-matched IDPs derived from the human proteome (biological hereafter) predicted to be strong clients (or excluders) and lacking any known interaction with FUS. We next sought to test the predicted recruitment of these IDPs (2 de novo, 2 biological) into FUS LC condensates in living cells.

We adapted a condensate assay to test predictions in which mCherry-tagged IDP sequences were co-expressed alongside CFP-tagged FUS LC fused to the DNA-binding LacI domain. As previously described^56^, the LacI domain localizes FUS LC to an array of LacO DNA binding sites integrated into the genome of U2OS cells, thus enabling simultaneous imaging of condensation as well as recruitment/exclusion of IDPs (see SI Note 4, Fig. 4B) at a single locus. Upon expression, we find that cells typically exhibited a large micron-scale FUS LC condensate in the nucleus (Fig 4C).

Consistent with predictions, we find that de novo clients partition into while de novo excluders are selectively depleted from the condensate (Fig 4C). To quantify protein enrichment across cells, we developed an image analysis pipeline (see SI Note 4) to automatically identify large, micron-scale FUS LC condensates and quantify IDP recruitment. For each of the 4 tested IDPs, we then averaged protein intensities in a 3 × 3 μm box centered around the condensate (Fig. 4D) and quantified the radial distribution of mCherry signal from the center of each FUS LC droplet (Fig. S10). Consistent with predictions, we found that FUS LC condensates were enriched for clients relative to the nuclear background and depleted for excluders. We observe that the extent of depletion for excluders is generally not as strong as that for recruitment (Fig. S10). Together, our results demonstrate our design framework enables programmable design of IDP interactions and guides their localization in living cells.

## Discussion

Intrinsically disordered proteins and protein regions (IDPs) are dynamic biomolecules that lack a stable fold and instead sample a wide ensemble of 3D conformations. This conformational plasticity underlies their key roles in cellular processes including binding, sensing, condensation, signaling, and localization, and are associated with pathological states when mutated or dysregulated. There is growing interest in designing de novo IDPs with tailored ensemble-derived properties, an area that is limited by the difficulty in approximating and inverting the dynamic sequence-ensemble-property relationships that characterize IDPs.

In this paper we develop a framework to flexibly and efficiently design IDPs by inverting physics-based models. Using this framework and validated through molecular simulations, we demonstrate that we can tailor ensemble dimensions i.e., the radius of gyration (Rg), for single IDPs and scalably generate proteome-scale de novo IDP libraries (Fig. 1). The modularity of our framework allows flexible incorporation of models and parameters beyond those described here, typically only requiring differentiable reimplementation. Promising avenues include incorporation of recent statistical sequence-based models^31,57,58^ and physics-based force fields^59,60^ with evolving accuracy.

Across biological organisms, IDPs often sense environmental cues^7,33,34^ through altered chain dimensions – a form of signal transduction where sensing, processing, and response are not embedded in different modules (proteins) but emerge from a collective physical process from a single module (IDP). Motivated by this, we program IDPs (Fig 2) that perform diverse computations such as threshold activation, bandpass response, and multi-input logic operations in response to changing environmental (temperature, salt, pH) and biological (phosphorylation, methylation) cues. In general, delineating how the physics underlying IDP ensembles constrains and enables molecular information processing and contrasting with other cellular computations that leverage collective^61–63^ as well as modular^36,64^ molecular networks are of future interest.

Multicomponent mixtures of IDPs often self-organize through phase transitions, including with other biomolecules, to form coexisting condensates. Biological condensates are characterized by rich emergent features that often shape their function - including formation of coexisting phases, layered morphologies, and activity-dependent remodeling. Inspired by this, we leverage our framework to rationally program multicomponent, multiphase condensates that exhibit diverse phase behavior, composition, layering, and RNA-dependent changes in architecture (Fig. 3). We then designed IDPs predicted to selectively partition in (client) or out (excluder) from condensates driven by the well-studied low-complexity domain of the FUS protein. Finally, we test and experimentally validate our predictions through localization assays of clients and excluders to FUS LC condensates in living cells (Fig 4). While we focus on localization, designing IDPs that rewire condensate phenotypes such as solubility, composition, and material properties is an avenue towards selectively targeting aberrant aggregates that are associated with multiple pathological states and diseases^65–67^.

The interplay of machine learning approaches and physics-based models^1,11,13,23,68,69^ continues to expand the toolkit for flexible design of biomolecules. Combining recent advances in design of structural folds^9,10,70–72^ with disordered protein design will enable generative modeling of multi-domain proteins. Further, designing emergent dynamic behaviors, including kinetics of aggregation^73^, transient occupation of functional conformer states^1,74^, and reactive condensates^75,76^ remains an important subsequent challenge. Together, the ability to predict and control sequence, conformational ensemble, and emergent properties represents an exciting frontier for biomolecular design.

## Limitations

The coarse-grained theories we employ capture effective properties such as pairwise interaction strengths but, by construction, abstract away finer, residue-specific geometries and rare conformation statistics that may shape IDP properties in vivo. Hence, our approach is unable to generalize to arbitrary simulation-derived quantities. Second, since existing physics-based models only capture equilibrium properties, nonequilibrium features that may be relevant to cellular function cannot be explicitly designed. Third, the effect of cues are captured approximately: changes in salt concentration are treated at the level of screened electrostatics and do not capture ion-specific effects, and post-translational modifications (phosphorylation and methylation) are represented through effective changes in residue charge rather than the detailed chemistry they introduce. Finally, although our framework is model-agnostic, it should be noted that the accuracy of any design, including the ones presented in this work, is ultimately dependent on the accuracy of the model it was designed over, meaning this framework will benefit from the development of increasingly accurate predictors of IDP behavior.

## Data and code availability

All protein sequences, figure-associated data, and code for the design presented in this work are available along with manuscript and/or github.

Link to github: https://github.com/shrinivaslab/2026_tyagi_boodry_IDP_design

## Acknowledgments

We thank Ryan Krueger and members of the Shrinivas lab for helpful discussions on this manuscript. We thank Benjamin Sabari for providing the pcDNA3.1-mCherry and CFP-LacI-FUS LC plasmids, as well as the U2OS 2-6-3 (LacO array) cells. The cell line was originally created by the lab of David Spector. J.B. acknowledges support in part by the National Institutes of Health Training Grant (5T32GM140995) through Northwestern University’s Molecular Biophysics Training Program. N.T. acknowledges support from the CSB Fellows and Walder Bridge Funds. K.S. acknowledges support in part by grants from the NSF (DMS 2235451) and Simons Foundation (MPS-NITMB-00005320) to the NSF-Simons National Institute for Theory and Mathematics in Biology (NITMB). W.T.S. acknowledges support from the NIH (R00GM149757). J.B., N.T., K.S., W.T.S., and V.C. acknowledge the support of startup funds from Northwestern University. This research was supported in part through the computational resources and staff contributions provided for the Quest high performance computing facility at Northwestern University.

