## Supporting Information for "Physics-guided design of intrinsically disordered proteins"

**This PDF file includes:**

SI Notes 1-4
SI References
SI Figures S1-S10

#### SI Note 1: Sequence design

##### Training and related hyper-parameters

We performed protein sequence optimization in python+jax using the gradient optimizer Optax with the adaptive moment estimation method (Adam) and employed a learning rate of 0.1. All single-chain optimizations were run for a total of 500 steps, while optimizations considering inter-IDP interactions, such as those reported in Figures 3 and 4, were run for 2500 steps.

##### Note on probabilistic sequence parameterization

We use a probabilistic sequence representation (see Krueger and Shrinivas[1] for details) to leverage continuous gradient-based updates, described briefly here. Consider a system of *n* particles (for example representing sequence length), each of which may be ascribed one of *m* identities (for example 1 out of 20 amino acid labels). Let $\pi \in \mathbb{R}^{n \times m}$ denote a matrix of particle identity probabilities, such that $\pi_{ij}$ represents the probability that position *i* in the sequence has identity *j.* Note that for any discrete sequence, $\pi$ is one-hot, meaning each position is represented by a vector of length *m* with all entries but one being 0. In practice, optimizations update an unnormalized logits matrix ($\lambda \in \mathbb{R}^{n \times m}$) that must be normalized to return $\pi$, i.e. $\pi = softmax(\lambda)$. Because the final desired output of each optimization procedure is a discrete sequence, we include a temperature $\tau$ to the normalization as follows: $\pi = softmax(\lambda/\tau)$. Lower temperatures return more discretized sequences and vice-versa. We find that a linear annealing with $\tau_{start} = 1.0$ and $\tau_{end} = 0.001$ worked well for single-chain designs (Figures 1 and 2), while we used $\tau_{start} = 1.0$ and $\tau_{end} = 0.01$ for optimizations with inter-IDP interactions (Figures 3 and 4).

##### Design criteria for each task

###### Single-chain Rg

We optimized sequences of a fixed length (N = 50) using the dimer model across a range of self-interaction parameters, χ, from -0.6 to 1 at intervals of 0.1 (with 17 points in total). This range represents the achievable limits for this fixed sequence length and choice of force field (calvados)[2,3]. For each value of χ, we generated a set of 1000 sequences and chose 25 at random to simulate using molecular dynamics to measure the radius of gyration ($R_g$, Figure 1B). Using this dynamic range for reference, we span self-interaction values for other models i.e., FINCHES and RPA. Specifically, we chose a range of linearly-spaced points from $-35 < \varepsilon < 35$ for FINCHES (11 points total) and logarithmically-spaced points from $0.3 < \chi_{RPA} < 10$ (13 points total) for RPA. As before, we then generated 1000 sequences per value and randomly chose 25 to simulate using molecular dynamics. These data are shown in Figures 1B, right and S3 for FINCHES with the calvados force field and RPA with the MPIPI-GG force field, respectively.

#### Stimuli-Responsive IDPs

We used the dimer model + CALVADOS to design IDPs that either expand or contract in response to a change in one external stimulus (temperature, salt concentration, pH, phosphorylation or methylation state). The ensemble dimensions are effectively encoded through requiring χ to be values that correspond to contracted or expanded chains in response to specified environmental cues. Motivated by Flory-Huggins theory[4], as well as the relationship between range of χ and chain dimension in Fig. 1B, we require that contracted chains (or expanded chains) have χ that is $> 0.5 + 1/\sqrt{N}$ (or $< 0.5 - 1/\sqrt{N}$). Single dimensional sensors had χ values calculated at two conditions: physiological and modified (Fig 2A, SI Table 1) as below:

| Condition | Physiological State | Modified State |
|---|---|---|
| Temperature | 300 K | 360 K |
| Salt Concentration | 150 mM | 450 mM |
| pH | 7.4 | 8 |
| Phosphorylation | $S^0$ | $S^{2-}$ |
| Methylation | $R^{1+}$ | $R^0$ |

For all sensors, the condition chosen as the stimulus was modified between the physiological and modified states, while other conditions were kept at the physiological values. See details in Parameterizing Cues to understand how each condition is incorporated into the physics framework. We used this methodology to design both the single-dimensional and two-dimensional sensors (Figures 2A and 2D). All of these sequences have sequence length 50, with the phosphorylation and methylation sensors having 6 modifiable sites.

The threshold-based sensors in Figure 2B were designed using the same cutoffs for χ and physiological/modified condition states, though these sequences had length 100 and contained 12 sites for phosphorylation/methylation. Threshold design is incorporated into our framework by additionally specifying χ values at intermediate points between the physiological and modified states such that the sequence's χ value changes appropriately. The bandpass sensor presented in Figure 2C specifies additional points with changing stimuli This sequence is of length 50 with 6 phosphorylation sites and begin and end at χ = 0.5, contracting in between.

#### Programming condensates with multiple IDPs

We optimized the interaction network of multiple IDPs for desired condensate behavior. In general, this is a many-to-one map, so we aim for conditions that generate our design of choice. All sequences designed here have length N = 50. The Coexistence phenotype (Figure 3B) requires 2 coexisting condensed phases, each with 2 species that condense. Hence, we impose strong self-interactions for all species ($\chi_{ii}$ >= 0.8) favorable interactions for species that co-condense ($\chi_{ij}$ <= -0.3), and unfavorable interactions for species that don't mix ($\chi_{ij}$ >= -0.1). To generate 3 layers (Figure 3C), we progressively ordered the self-interaction strengths (and thus surface tensions) in our design objective while ensuring all species still remained condensed in distinct phases by minimizing cross-species interactions. Specifically, the outermost sequence satisfies $\chi_{11}$ >= 0.65, the middle one $\chi_{22}$ >= 0.75, and the innermost $\chi_{33}$ >= 0.9, while cross-interaction parameters were specified as $\chi_{12}$ >= -0.10, $\chi_{13}$ >= -0.20, and $\chi_{23}$ >= -0.25. For the Composition phenotype (Figure 3D), our goal was to create a condensate with 2 species that did not recruit 2 other species that remain soluble. Condensed species interact strongly (self-interaction $\chi_{ii}$ >= 0.8 and cross-interaction $\chi_{ij}$ <= -0.3) while soluble species have disfavorable self-interaction $\chi_{ii}$ <= 0.0 and cross-interaction $\chi_{ij}$ >= 0.5. Interactions between condensed-species and soluble-species are specified to be $\chi_{ij}$ >= 0.01. For RNA-dependent remodeling (Fig 3E), our goal was to design a condensate rich in 2 IDPs, that upon addition of RNA, would selectively enrich RNA and deplete 1 of the IDP species. Towards this, the two IDPs were designed to satisfy $\chi_{ii}$ >= 0.6 for self-interaction and $\chi_{ij}$ <= -0.2 for inter-IDP interactions. For IDP-RNA interactions, the chain that mixes with RNA must satisfy $\chi_{i,RNA}$ <= -0.35 while the other must satisfy $\chi_{i,RNA}$ >= -0.22. RNA in this system as poly uracil (polyU) with a length of 50 nucleotides.

#### Clients and Excluders

For design of clients and excluders, we optimized the interaction between designed IDP and FUS LC domain ($\chi_{i,FUS}$). We first analyzed biological IDPs derived from human proteins with length $130 \leq N \leq 150$ and annotated as localizing to the nucleus (sources - UNIPROT[5]). Since $\chi$ is a length-normalized quantity (see Physics Models), we calculated $N_i\chi_{i,FUS}$ values for each IDP in this subset of the human IDPome to remove any length dependence (see Fig. S9, left for full $N_i\chi_{i,FUS}$ distribution). We selected proteins from this distribution of $N_i\chi_{i,FUS}$, choosing Zinc finger protein 69 homolog B[6] (Uniprot ID: Q9UJL9), residues 131-267 as the biological client ($N_i\chi_{i,FUS}$ = -6.14) and Zinc finger protein 316[7] (Uniprot ID: A6NFI3), residues 1-150 as the biological excluder ($N_i\chi_{i,FUS}$ = 2.92). We then optimized de novo client and excluder sequences to match these biological examples, each with length $N_i = 140$ with the client having a specified $N_i\chi_{i,FUS}$ value of -6.0 and the excluder 3.0.

#### SI Note 2: Physics models

We provide a high-level overview of physics-based models used in our paper and expanded details are available in the original publications[2,8,9].

##### Dimer Model

The dimer model aims to estimate the second virial coefficient (B) from IDP sequence identity by considering all nonbonded interactions between all pairs of adjacent amino acids in the protein sequence according to the following equation:

$$B^{DP}(T) := \sum_{n=1}^{N-1}\sum_{m=1}^{N-1} \nu^{DP}_{d_{n,n+1}d_{m,m+1}}(T)$$

Where $\nu$ refers to the virial coefficient for a given pair of amino acid dimers, calculated as:

$$\nu^{DP}_{d_{n,n+1}d_{m,m+1}}(T) := 2\pi\int_0^\infty drr^2[1 - e^{-\beta U^{DP}_{d_{n,n+1}d_{m,m+1}}(r)}]$$

With the explicit energy dependence of $U$ on dimer identity given by:

$$U^{DP}_{d_{n,n+1}d_{m,m+1}} = U_{a_n a_m} + U_{a_{n+1}a_m} + U_{a_n a_{m+1}} + U_{a_{n+1}a_{m+1}}$$

Where each substituent term, $U_{i,j}$, includes energetic contributions between amino acids *i* and *j* arising from all nonbonded interactions. As defined in this work, the length-normalized self-interaction parameter $\chi_{ii}$ is given by the following relationship derived from the virial expansion for homopolymers in Flory-Huggins polymer theory[4]:

$$\chi_{ii} = 0.5 - \frac{B_{ii}}{N_i^2 l^3}$$

Where $l$ refers to the lattice constant in the original Flory-Huggins formulation. Adachi and Kawaguchi[2] found a suitable value for $l = 3.6l_b$, where $l_b = 0.38\,nm$ is the bond length between amino acids which we employ in this work. The cross-interaction parameter $\chi_{ij}$ is defined as:

$$\chi_{ij} = \frac{B_{ij}}{N_i N_j l^3}$$

##### Random Phase Approximation Model

The RPA model specifically considers contributions from three primary sources to predict net interaction between protein sequences. First is a mean-field approximation related to an effective Flory-Huggins $\chi$ parameter determined from pairwise residue-residue interactions. Second is a mean-field contribution from screened electrostatic interactions across the entire chain length dependent on the average residue charge of each chain. Finally, the RPA-specific contribution emerges from electrostatic interactions as a function of spatial charge distribution within the chain itself.

#### FINCHES Model

The FINCHES model incorporates specific sequence-level motifs into the prediction of interaction or repulsion between sequences. First, FINCHES calculates a pairwise interaction matrix between all residues in the two sequences treating them as “bags of monomers”. In addition, FINCHES analyzes local sequence context for all residues in both proteins, focusing on charged and aliphatic interactions. Local charged interactions are accounted for by analyzing the $i + 1$ and $i - 1$ charges around a charged residue and down-weighting like-charge interactions, effectively reducing the repulsion associated with groups of like-charged residues. Local hydrophobic effects arising from clustered aliphatic residues are accounted for by considering continuous runs of 2 and 3 (or more) aliphatic residues, upweighting aliphatic-aliphatic interactions for these clusters by $1.5\times$ and $3.0\times$ for residues embedded in runs of 2 and 3 (or more) aliphatic residues, respectively. Finally, sequence-level interaction values $\epsilon$ values are obtained by summing this reweighted interaction matrix. Note that these values are extensive quantities.

#### Model benchmarking

To ensure agreement between the original and differentiable implementations of each physics model, we began by generating 10,000 random amino acid sequences of length 50 as a benchmarking dataset. For each force field available in the original model implementations, we calculated the predicted interaction parameter value ($B^{DP}$, $\epsilon$, $\chi_{RPA}$) for all 10,000 sequences and compared with the differentiable implementation. This analysis is shown in Figure S1, notably demonstrating a 1-1 correspondence between both model instantiations.

#### Parameterizing cues

##### Temperature

Temperature-specific effects are incorporated into our prediction and design framework only through energetic calculations. Each physics model requires the computation of an integral with Boltzmann-weighted energies, thus temperatures are considered implicitly when calculating $e^{-\beta U}$, where $\beta = 1/k_B T$ is the inverse temperature.

##### Salt Concentration

Salt effects are considered in the calculation of the debye screening parameter, $\kappa$, which affects the magnitude of coulombic energy contributions. The equation we used to determine this $\kappa$ value as a function of salt concentration is given by the following:

$$\kappa = \kappa_0 \sqrt{\frac{C_s}{0.150}}$$

Where $\kappa_0$ refers to the force field specific debye screening factor at physiological (150 mM) salt concentrations, and $C_s$ is the salt concentration in M.

### pH

Unless otherwise noted, the charges on amino acid residues for all designs presented in this work correspond to force field definitions. In the cases corresponding to the design of IDPs that respond to pH as a stimulus (table S1), residue charges were modified using the Henderson-Hasselbalch equation, given below:

$$q = \frac{\pm 1}{1+10^{\pm(pH-pKa)}}$$

##### Phosphorylation and Methylation

Phosphorylation and methylation act within our optimization procedure simultaneously as conditions and design constraints. These stimuli are treated within our framework through modified residues that mimic these effects as follows. In the unmodified state, these sites are one-hot encoded to represent either serine or arginine (e.g. $S^0$ and $R^{1+}$), while in the 'modified' state these sites switch to a one-hot encoding of the corresponding modified residues. We represent phosphorylation by using the unmodified serine force field parameters for non-coulombic nonbonded interactions, but including a charge of 2- on the residue ($S^{2-}$) as has been done previously[10], while methylated sites are represented by an uncharged arginine residue ($R^0$).

#### Molecular dynamics simulations

We used the CALVADOS[3,11] package to run MD simulations on all designed sequences using OpenMM[12] and the CALVADOS force field. Model formulation details, including force field parameterization, are given in the original paper, while we provide a brief overview of simulation setup here for both single-chain and slab simulations. All simulations were run using a Langevin integrator with a 10 fs timestep and periodic boundary conditions.

##### Single Chain Simulations

We ran single chain IDP simulations using a cubic simulation box with side length of 50 nm. Within the CALVADOS simulation package, we specified a centre topology with a single molecule. For other input conditions, salt concentration, pH, phosphorylation, and methylation are incorporated identically to the formulation in our main design loop (see Parameterizing Cues). Temperature effects are included implicitly as an argument to the Langevin integrator. Single chain simulations are run for a total time of 100 ns, with 1 ns of equilibration which is discarded when performing analysis. The CALVADOS package additionally includes functionality to automatically calculate radii of gyration as simulations are performed, which we use for our analysis in Figures 1 and 2.

##### Slab Simulations

Slab simulations are run in an elongated simulation box of size $15\,nm \times 15\,nm \times 150\,nm$, enabling the study of dense and dilute protein phases in condensed-phase systems. Initial chain placements are determined automatically within the CALVADOS simulation package by specifying a slab topology. These simulations are run for 1 μs with 10 ns of equilibration. IDP designs presented in Figure 3 of this work contain 100 copies of each de novo protein sequence, with the RNA-dependent design additionally containing 60 copies of length 50 polyU RNA. Slab simulations for clients and excluders presented in Figure 4 contain 100 copies of the FUS LC domain with a single copy of either the client or excluder. We performed density profile analysis of slab simulations directly from the output trajectory files, averaging over 10,000 total frames with 0.1 ns spacing between frames, to determine the spatial distribution of various IDPs along the extended simulation box axis.

#### SI Note 3: Sequence analysis

##### Dataset construction

We compared sequence features from de novo designed IDPs with human derived IDPs[5]. From the human proteome, we first calculated the per-residue disorder score for each protein in the dataset using Metapredict v3, defining disordered regions as continuous sequences of at least 30 amino acids with per-residue disorder above 80%. We subset this group focusing on sequences with length $30 < N < 70$, representing 8,521 biological sequences. High dimensional analysis proceeded with two different datasets: the first contains all human IDPs as defined above and de novo IDPs designed (using the dimer and FINCHES models) to match the distribution of $\chi$ (or $\epsilon$ for FINCHES) present in the human IDPome. The second dataset contained the same subset of the human IDPome, but sequences were designed with a uniform interaction parameter distribution across the dynamic range of self-interactions using the dimer and FINCHES models.

##### Clustering and dimensionality reduction

We generated encodings from sequences to perform high-dimensional clustering following 2 different steps and our key results hold true both ways. The first is a composition-based encoding where each sequence is represented by a dimer frequency vector. Each entry represents the percentage of the sequence composition represented by each amino acid dimer (AG, GG, etc.). Second, we use tokenized sequence representations generated by the ESM2 protein language model[13]. Regardless of sequence encoding, we first perform principal component analysis (PCA) with the scikit-learn python package to reduce the dimensionality of the dataset before proceeding with clustering and visualization[14]. We generated embeddings with the UMAP package from umap-learn using these PCA-reduced datasets for use in visualization, with relevant hyperparameters n_neighbors = 15 and min_dist = 0.1[15]. UMAP arbitrarily distorts distances between points on the final produced visualization[15–17] and should be used as a guide. For clustering, we quantified sequence similarity through unsupervised K-Means clustering with the scikit-learn package on the PCA-reduced datasets. We determined the ideal number of clusters ($k$) through a cluster-stability based metric introduced by Asa-hur et al[18]. If multiple values of $k$ produced quantitatively similar cluster-stability results, we chose the largest number of clusters among these for a maximally-informative clustering. For this $k$ value, we analyzed composition of clusters based on where the sequences came from (IDPome, designed with dimer model, designed with FINCHES).

#### SI Note 4: Experimental design and assays

##### Plasmid design

The pcDNA3.1-mCherry and CFP-LacI-FUS LC plasmids were kindly provided by Benjamin Sabari[19]. The LC domain of FUS comprises amino acids 2-214. The pcDNA3.1-mCherry plasmid contained SV40 nuclear localization signals (NLSs) flanking mCherry, ensuring nuclear localization of the IDPs. Protein-coding sequences of interest were codon optimized for expression in mammalian cells (*Homo sapiens*) and synthesized as gBlock Gene Fragments (Integrated DNA Technologies). The coding sequences of interest were amplified by PCR and individually cloned into pcDNA3.1-mCherry backbone using Gibson Assembly Master Mix (New England Biolabs E2611S).

| Protein | Amino Acid Sequence | DNA sequence (5’ → 3’) | Fwd primer (5’ → 3’) | Rev primer (5’ → 3’) |
|---|---|---|---|---|
| client_bio_137/Q9UJL9_4 (Len = 137) | EPWLMERDISGVPSSDLKSKTKTKESALQNDISWEELHCGLMMERFTKGSSMYSTLGRISKCNKLESQQENQRMGKGQIPLMCKKTFTQERGQESNRFEKRINVKSEVMPGPIGLPRKRDRKYDTPGKRSRYNIDLV | GAGCCCTGGCTGATGGAGCGCGACATCAGCGGCGTGCCCTCTAGCGACCTGAAGTCTAAGACTAAGACTAAGGAGAGCGCCCTGCAGAACGATATTTCATGGGAGGAGCTGCACTGCGGCCTGATGATGGAGAGATTCACCAAGGGCTCCAGCATGTACAGTACACTGGGAAGAATCTCCAAGTGTAATAAACTGGAGAGTCAGCAGGAGAATCAGCGCATGGGCAAGGGCCAGATCCCTCTGATGTGCAAGAAGACCTTCACCCAGGAGAGGGGCCAGGAGAGTAATAGGTTCGAGAAGAGAATCAACGTGAAGTCTGAAGTGATGCCCGGCCCCATCGGCCTCCCAAGGAAGAGGGATCGGAAGTACGATACCCCCGGCAAGAGGAGCAGGTATAATATCGACCTGGTG | tgcgggttctgccgcaggtg*gatcc*gagccctggctgatggag | ccctctagagacggcgctcg*aattc*ttacaccaggtcgatattatacctgc |
| client_denovo_140 | CGNIPHSNCHSGMHSSNTHNITVMHNPTNHSNCTIMGHNKPNSVCASNLTISHCICTCICNHKNIQNHNHPQHNQPKKHNHCHHKHPGHNNHHQNHSNLCNHHPDPTQHCCQNTQIQNNGNINNNICMVNQPQHHPTSGI | TGCGGCAACATCCCACACTCCAACTGCCACTCTGGCATGCATAGCTCCAACACCCACAACATTACAGTGATGCACAATCCTACAAATCACTCTAATTGCACTATCATGGGGCACAACAAGCCAAATAGCGTGTGTGCCTCCAACCTGACCATCTCCCATTGCATCTGCACATGCATTTGCAACCACAAGAACATCCAGAACCACAACCACCCCCAGCACAATCAGCCTAAGAAGCACAACCATTGCCACCACAAGCATCCCGGGCACAATAACCACCACCAG | tgcgggttctgccgcaggtg*gatcc*tgcggcaacatcccacac | ccctctagagacggcgctcg*aattc*ttagatgccagaggtagggtg |

| | | | | |
|---|---|---|---|---|
| | | AACCACAGCAACCTGTGCAAC<br>CACCACCCCGACCCCACACAG<br>CACTGCTGTCAGAACACCCAG<br>ATTCAGAATAACGGCAATATTAA<br>CAACAACATTTGCATGGTGAAC<br>CAGCCCCAGCACCACCCTACC<br>TCTGGCATC | | |
| exclud<br>er_bio<br>_149/<br>A6NFI<br>3_1<br>(Len =<br>149) | MAALHTTPDSPA<br>AQLERAEDGSEC<br>DPDQEEEEEEEE<br>KGEEVQEVEEEE<br>EEIVVEEEEEGVA<br>EVVQDAQVEAVA<br>EVEVEADVEEED<br>VKEVLAEEECPAL<br>GTQERLSRGGDA<br>KSPVLQEKGLQA<br>SRAPATPRDEDLE<br>EEEEEEEDEDED<br>DL | ATGGCAGCACTCCATACAACACCT<br>GATTCCCCTGCAGCCCAGCTGGA<br>GAGAGCCGAGGATGGCAGCGAGT<br>GCGATCCTGACCAGGAGGAGGAG<br>GAGGAGGAAGAAGAGAAAGGGGA<br>AGAAGTGCAGGAAGTGGAGGAGG<br>AAGAAGAAGAAATTGTTGTGGAAG<br>AGGAAGAGGAAGGCGTCGCAGAG<br>GTTGTGCAGGACGCCCAGGTCGA<br>GGCAGTGGCCGAAGTGGAGGTGG<br>AAGCCGACGTGGAGGAGGAGGAC<br>GTGAAGGAGGTGCTGGCCGAAGA<br>GGAATGCCCTGCCCTGGGAACCC<br>AGGAGAGGCTGTCCAGGGGCGG<br>GGATGCTAAGTCCCCTGTTCTGCA<br>GGAAAAGGGACTGCAGGCTAGCA<br>GAGCCCCAGCCACACCAAGGGAT<br>GAAGACCTGGAGGAGGAGGAAGA<br>GGAAGAAGAGGACGAGGATGAAG<br>ACGACCTG | tgcgggttctgcc<br>gcaggtg*gatc*<br>*c*atggcagcact<br>ccatacaac | ccctctagaga<br>cggcgctcg*aa*<br>*ttc*ttacaggtcg<br>tcttcatcc |
| exclud<br>er_de<br>novo_<br>140 | VVKDAVAPVPVVE<br>DDAAVVADATDKA<br>DVVEDKTAPKPE<br>VEVDVVDVDVAV<br>EVDKADVEDVAA<br>DQDDEVVEVAVV<br>DAKSKTDTDDKP<br>EADDEDKAAAKD<br>VVVPAKVVDAPK<br>AKPVKAVKTPPVV<br>VVDVVVPEAAVV<br>VAAVG | GTGGTGAAGGATGCCGTGGCGCC<br>TGTGCCCGTTGTGGAAGATGATGC<br>CGCCGTGGTGGCCGATGCCACTG<br>ACAAAGCTGATGTTGTGGAGGATA<br>AGACTGCACCAAAGCCTGAGGTG<br>GAAGTCGACGTGGTGGATGTGGA<br>CGTGGCCGTTGAAGTGGACAAGG<br>CCGATGTGGAGGACGTGGCCGCC<br>GATCAGGACGACGAGGTGGTGGA<br>GGTGGCCGTGGTGGATGCCAAGT<br>CCAAGACCGATACCGATGATAAGC<br>CCGAAGCAGATGATGAAGACAAG<br>GCCGCTGCTAAAGATGTGGTGGTC<br>CCCGCTAAGGTTGTGGATGCCCCA<br>AAAGCTAAGCCCGTGAAGGCTGTT<br>AAGACACCTCCCGTGGTCGTGGT<br>CGATGTGGTGGTGCCCGAGGCTG<br>CCGTGGTGGT[20]GCTGCAGTGGGA | tgcgggttctgcc<br>gcaggtg*gatc*<br>*c*gtggtgaagg<br>atgccgtg | ccctctagaga<br>cggcgctcg*aa*<br>*ttc*ttatcccact<br>gcagcaac |

#### Cell Culture

U2OS 2-6-3 cells, containing an integrated array of LacO DNA repeats in the genome, were generated by the lab of David Spector[20] and were kindly provided by the lab of Benjamin Sabari. U2OS 2-6-3 cells were cultured in Dulbecco's modified Eagle's medium (DMEM), supplemented with 10% fetal bovine serum (FBS), 1% penicillin-streptomycin, and 1% GlutaMAX Supplement (Fisher Scientific 35050061). Cells were maintained at 37°C in a humidified 5% $CO_2$ incubator.

#### Transfection

U2OS 2-6-3 cells were seeded onto acid washed glass coverslips in 12-well plates at 50% confluency and transfected the following day using Lipofectamine 3000 (Fisher Scientific L3000015) following manufacturer's suggested protocol. Cells were transfected with 1.5 µg pcDNA3.1-mCherry fused to the IDP of interest and 1.5 µg CFP-LacI-FUS LC per well. 24 hours post-transfection, cells were fixed with 4% paraformaldehyde, permeabilized with 0.5% Triton X-100 and stained with Hoechst (1:1000 dilution). Washes were performed with 1X PBS between each step. Coverslips were mounted onto glass slides using VECTASHIELD Vibrance® Antifade Mounting Medium and imaged as described below.

#### Microscopy

Imaging was performed on a Nikon ECLIPSE Ti2E inverted microscope system controlled by NIS-Elements AR (version 6.10.02). The system was equipped with an X-light V3 spinning disk confocal unit (CrestOptics) and images were acquired using a Nikon Lambda D oil immersion 100x objective. Samples were illuminated with a CELESTA Light Engine (Lumencor) using 365 nm (DAPI), 440 nm (CFP), and 561 nm (TRITC) excitation wavelengths. Images were captured using a Photometrics Kinetix sCMOS camera (Teledyne). Z-stacks were acquired with a step size of 0.2 µm. Identical laser power settings were used across conditions, and images were corrected to account for camera exposure time differences.

#### Image analysis

Multichannel images (DAPI for DNA, CFP for FUS LC, and mCherry for IDP) were analyzed as follows using a custom Python script. For each image, the full stack was read and a maximum-intensity projection (MIP) image was generated. Nuclear segmentation was performed on DAPI channel MIPs using Cellpose (v4.0.7)[21]. For each segmented nucleus, mean FUS LC intensity was measured from the exposure-normalized FUS LC image, and nuclei with mean FUS LC intensity >1000 were retained for downstream analysis and confirmed to also express IDP. For each retained nucleus, the original z stack was revisited to identify the z-plane with highest FUS LC signal in that nucleus and nucleus-specific 2D crops were extracted for each channel. These crops were used for all subsequent analyses.

From each crop, FUS LC condensates were detected using Otsu thresholding[22] and a minimum diameter of 0.5 um was enforced to primarily quantify LacO-associated large condensates. Condensate morphology and intensity descriptors were recorded and the largest qualifying condensate was selected per nucleus. For these condensates, radial intensity profiles for FUS LC and IDP signals were computed from the centroid to 2.0 µm with 0.1 µm bin width.

Condensates near cropped nuclear boundaries were excluded to avoid truncation artifacts. Per-nucleus radial profiles were grouped by condition to calculate condition-level mean profiles. Stacked images for each IDP were generated by extracting fixed 3.0 x 3.0 μm windows centered on the largest FUS LC condensate in each nucleus, and stacks were averaged to generate average fluorescence intensity images displayed in Fig. 4.

### Supplementary Figures

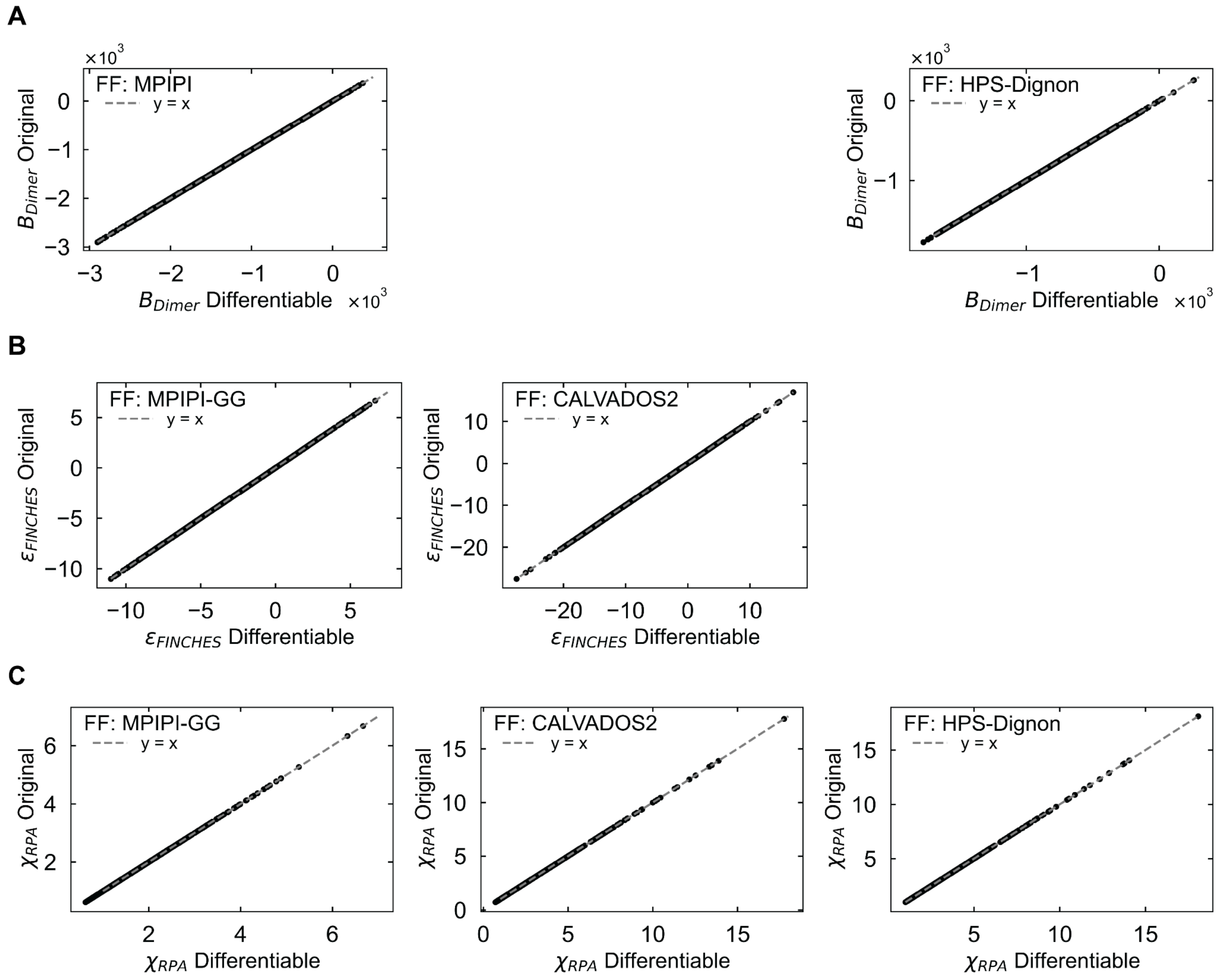


**Figure S1. Benchmarking**

(A-C). The predictions from the differentiable implementation of physics models (x-axis) are compared to the original implementations (y-axis) and each dot is an individual sequence while the dashed line is the y=x line. The force field of choice is listed inset and models considered are dimer (A), FINCHES (B) and RPA (C).

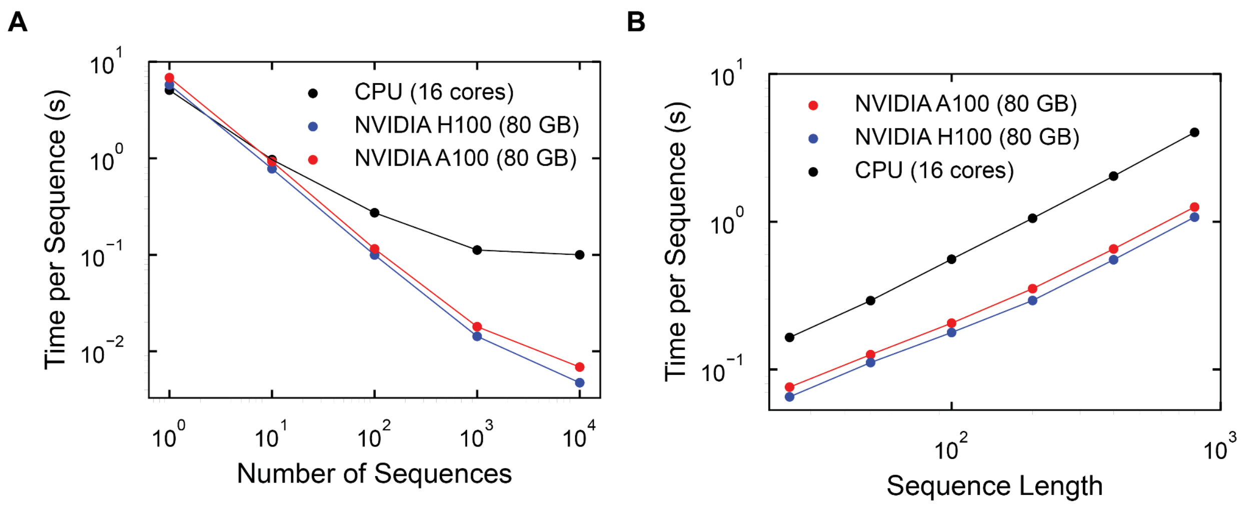


**Figure S2. Compute time scaling**

(A) Time required per-sequence (y-axis) for batch sequence optimization versus the number of sequences (x-axis) being optimized. (B) Time required per-sequence for batch (y-axis) sequence optimization versus sequence length (x-axis), where batch size is 100 sequences for all sequence lengths.

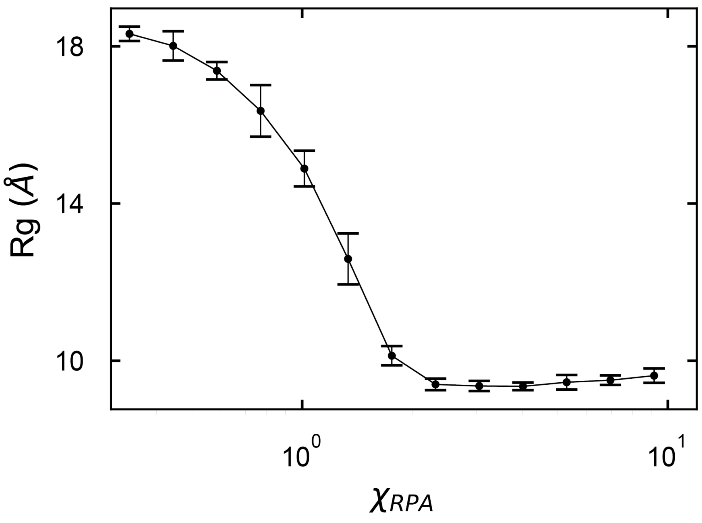


**Figure S3. RPA model comparison to simulations**
Simulation derived radius of gyration (Rg, y-axis) versus interaction parameter $\chi_{RPA}$ (x-axis) with the Mpipi-GG forcefield. Each dot is the mean of 25 sequences, error bars are the standard deviations, and the line is provided as a guide to the eye.

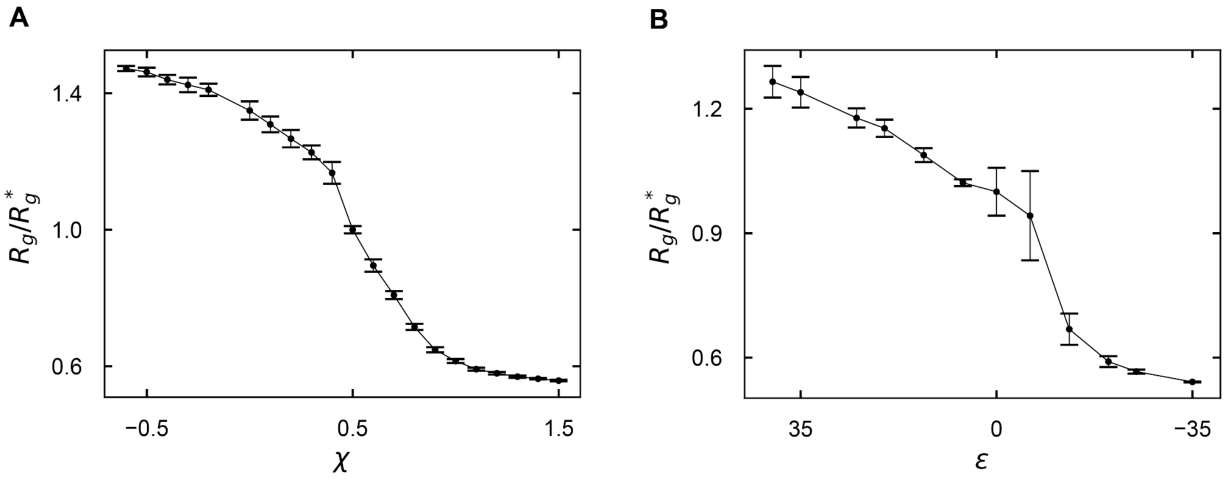


**Figure S4. Comparison of MPIPI-GG sequence optimization to simulation**
Simulation-predicted Rg values (y axis) versus self interaction values (x-axis) for dimer (A) and FINCHES (B) models.The Rg values are normalized to the average Rg* at χ=0.5 and ε=0. Each dot is the average over simulations of 25 independent designs, error bars are standard deviations, and the black line is a guide.

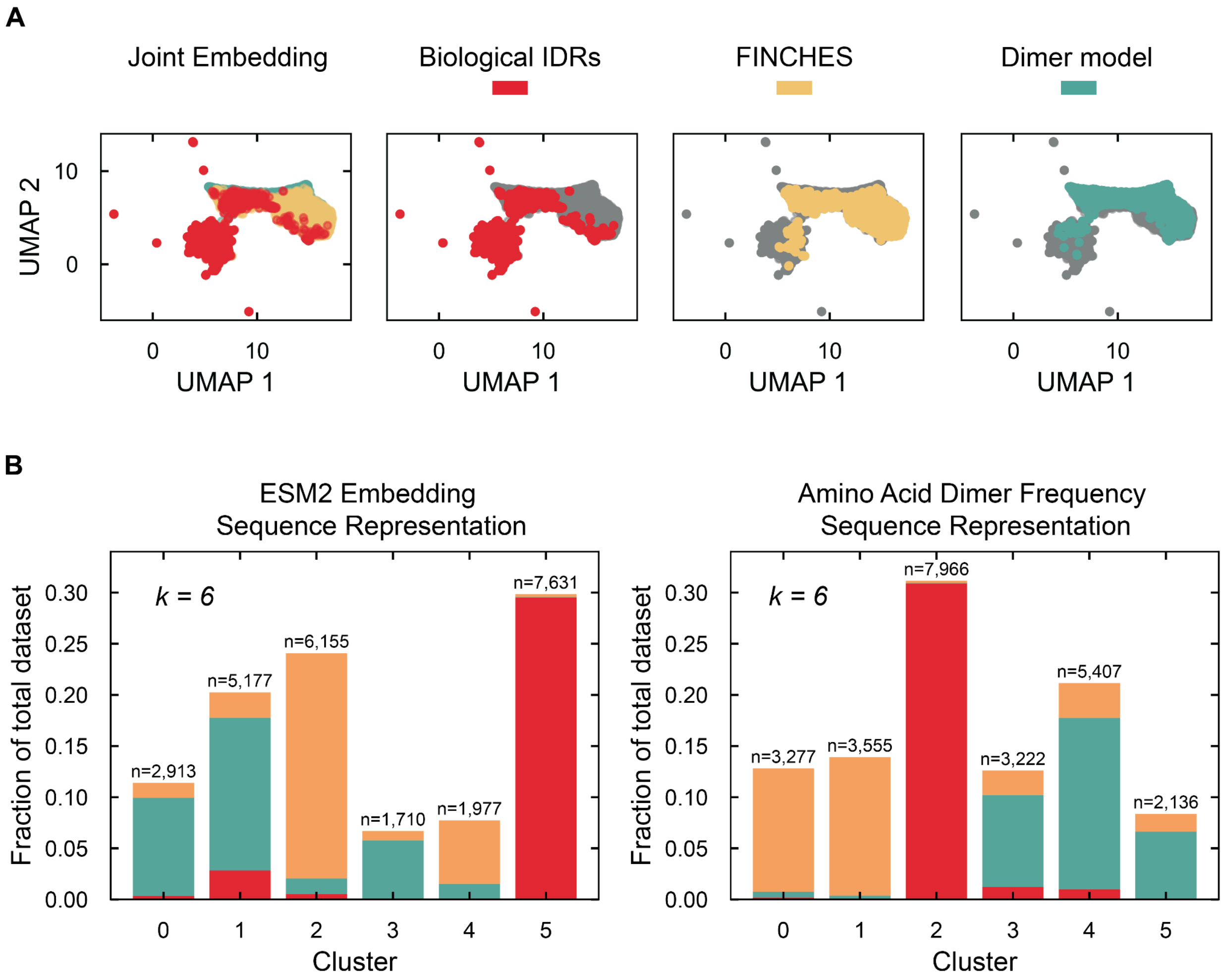


**Figure S5. Clustering of interaction-matched designed sequences with human IDPome**
(A) UMAP embedding generated using amino acid dimer frequency sequence representation. Biological IDPs are colored in red, de novo IDPs generated with FINCHES in yellow, and de novo IDPs generated with the dimer model in green. (B) Unsupervised K-Means clustering results arranged by composition (y-axis) of each cluster (x-axis) using ESM2 embeddings (left) and amino acid dimer frequency (right) to represent sequences. The color scheme is the same as (A).

See SI for extended details on analyses.

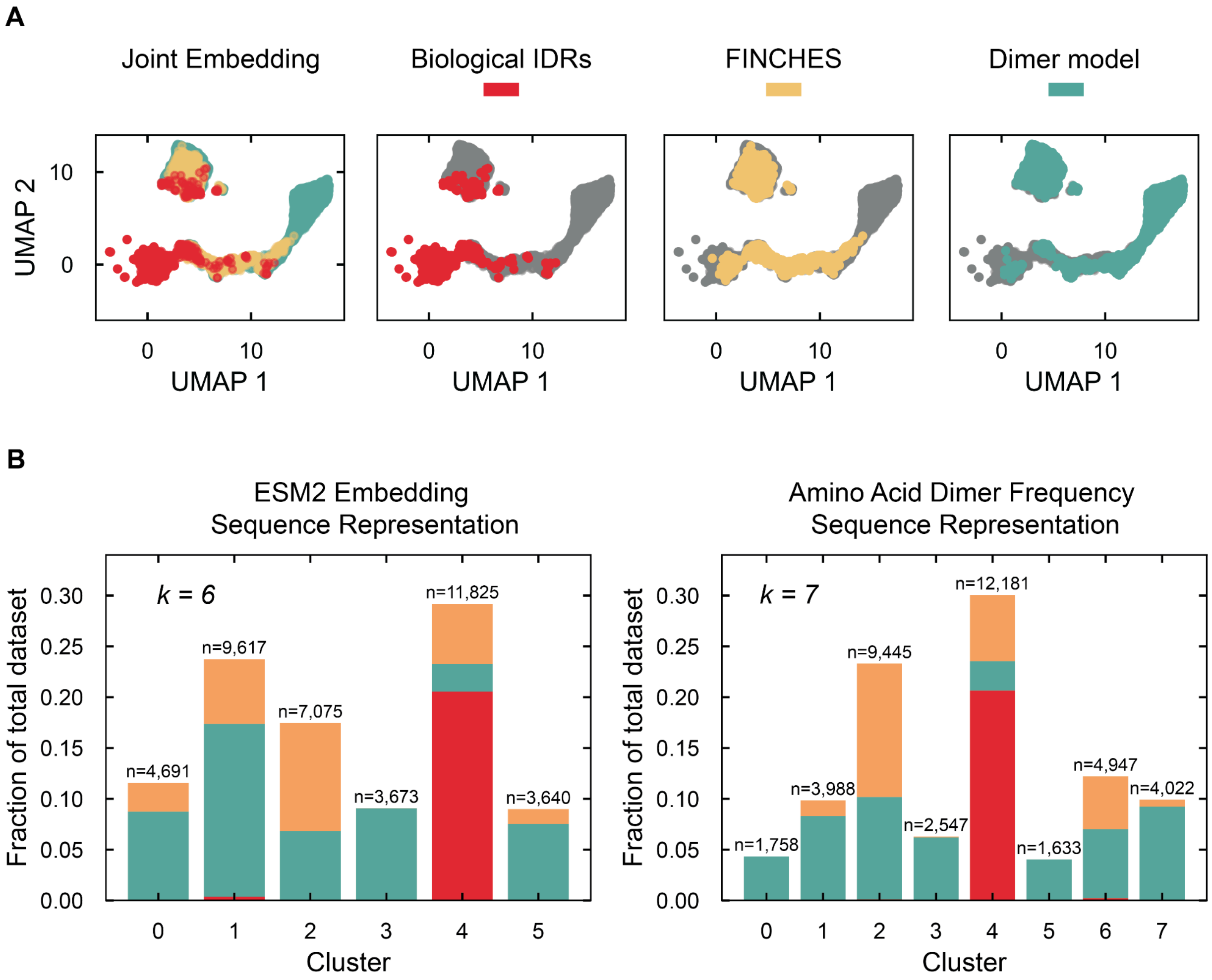


**Figure S6. Clustering of designed sequences with uniform interaction distribution with human IDPome**

(A) UMAP embedding generated using amino acid dimer frequency sequence representation. Biological IDPs are colored in red, de novo IDPs generated with FINCHES in yellow, and de novo IDPs generated with the dimer model in green. (B) Unsupervised K-Means clustering results arranged by composition (y-axis) of each cluster (x-axis) using ESM2 embeddings (left) and amino acid dimer frequency (right) to represent sequences. The color scheme is the same as (A).

See SI for extended details on analyses.

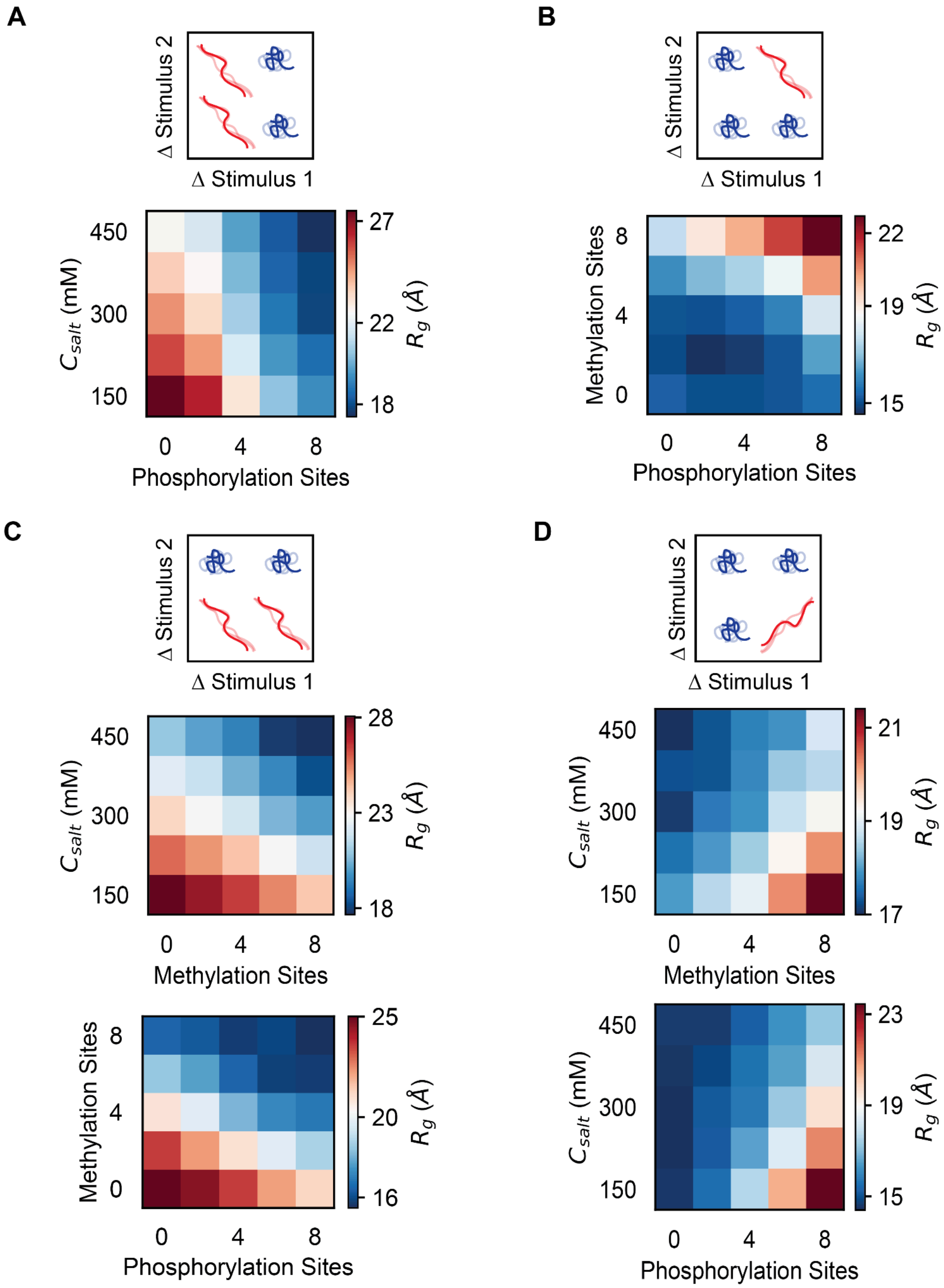


**Figure S7. 2-dimensional sensor designs**

(A-D) Schematic illustration of desired two-dimensional sensor design on top, with stimuli-dependent simulation derived Rg values below. Designs include: (A) A phosphorylation contractor that does not respond to salt concentration, (B) A methylation-phosphorylation AND gate, (C) two designs that contract in response to one stimulus and do not respond to another, and (D) two designs that are only expanded in one specific combination of stimuli. In all plots, the y and x-axis represent changing stimuli and the colors represent the Rg values with colorbar for scale.

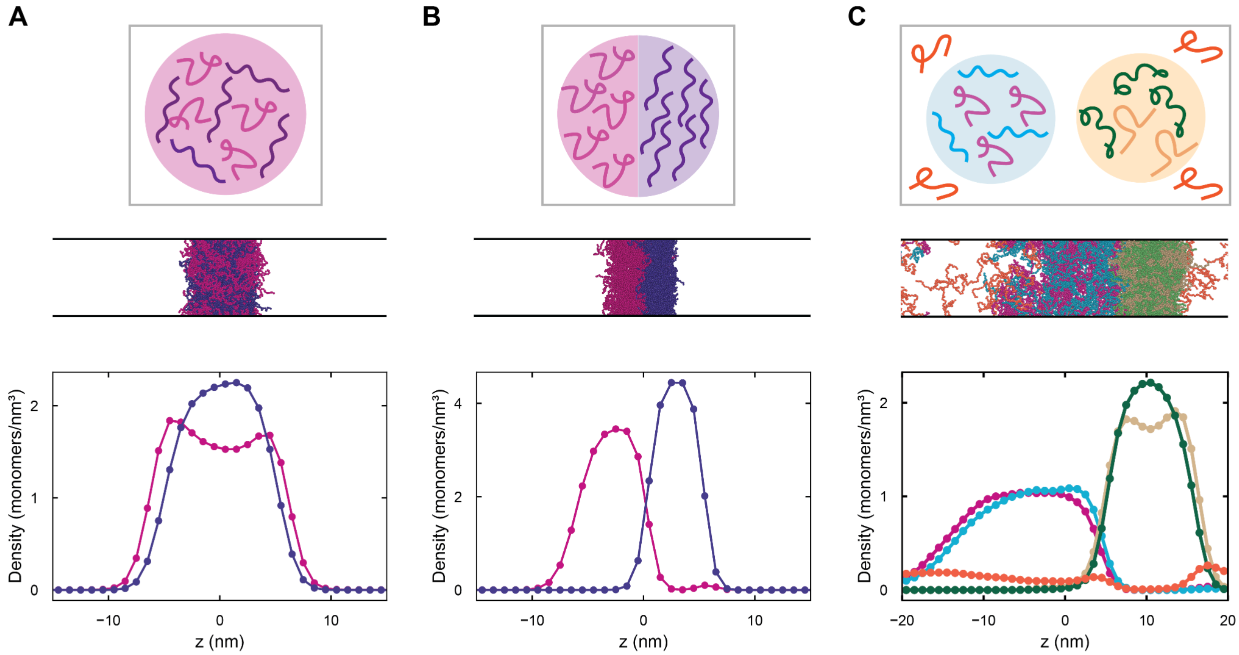


**Figure S8. Programmed multiphase behavior in multicomponent IDP mixtures**

(A-C) Schematic of desired phase behavior (top), representative snapshot from slab molecular dynamics simulation (middle), and quantified density profiles (y-axis) from simulations (bottom) is shown. Designs include a two-chain system that condenses and mixes into a single droplet (A), a two-chain system that condenses and forms separate droplets (B), and a five-chain system with two compositionally unique droplets and one chain in the dispersed phase (C).

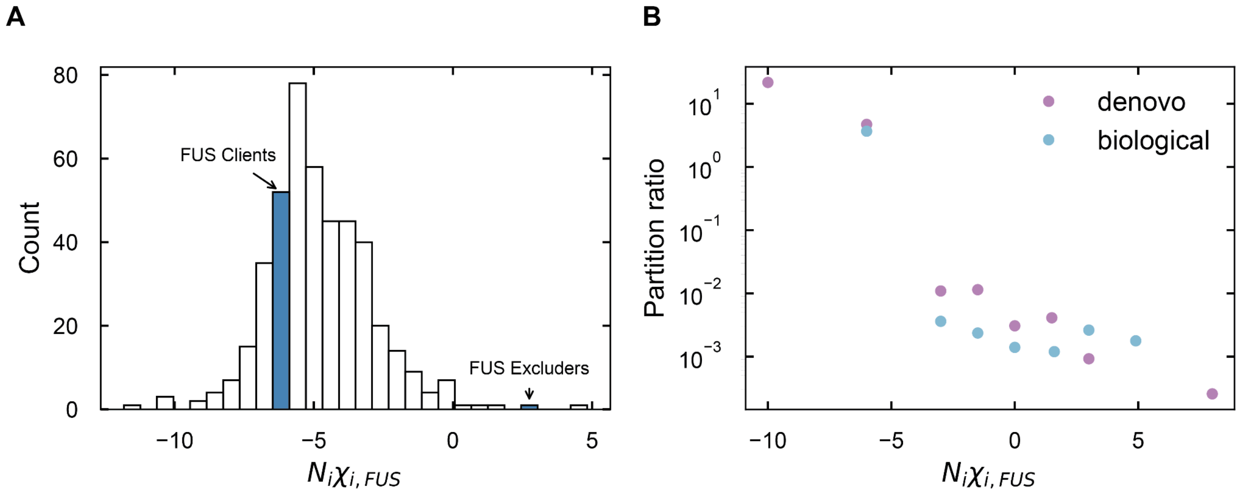


**Figure S9. FUS LC client and excluder identification**

(A) Histogram of $N_i\chi_{i,FUS}$ values (x-axis) for length-matched, nuclear IDPs in the human proteome. Bins corresponding to identified FUS LC clients and excluders are denoted. (B) Partition ratio (y-axis) versus $N_i\chi_{i,FUS}$ values (x-axis) for designed and biological IDPs obtained from slab simulation.

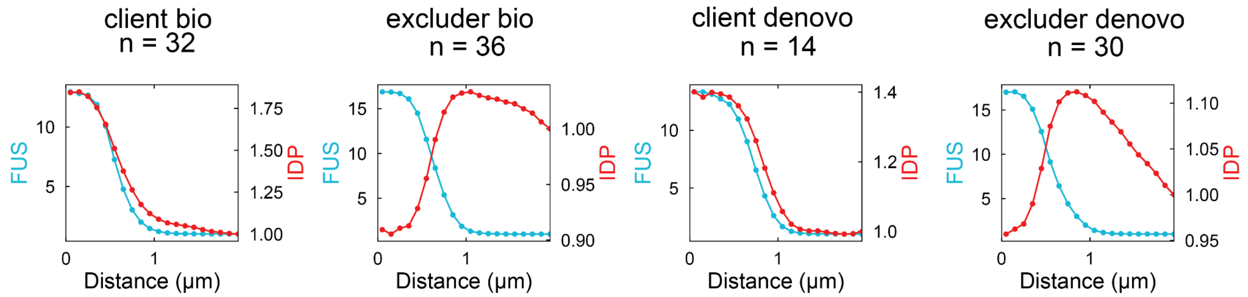


**Figure S10. Radial intensity of live cell assays**

Relative FUS LC (y-axis, left) and IDP (y-axis, right) intensities versus distance from the center of each FUS LC condensate (x-axis) and averaged across multiple condensates (specified in panel title). FUS LC and IDP intensities are normalized relative to the nuclear background.

**SI Table 1: 1-dimensional sensors**

| Stimulus | Response Type | $R_g^{low}$ (Å) | $R_g^{high}$(Å) |
|---|---|---|---|
| Temperature | Contractor | 16.0 | 15.9 |
| Salt | Contractor | 26.7 | 18.4 |
| pH | Contractor | 17.4 | 17.7 |
| Phosphorylation | Contractor | 22.5 | 15.9 |
| Methylation | Contractor | 24.0 | 19.6 |
| Temperature | Expander | 21.9 | 24.4 |
| Salt | Expander | 15.0 | 21.6 |
| pH | Expander | 19.8 | 21.4 |
| Phosphorylation | Expander | 16.5 | 22.9 |
| Methylation | Expander | 18.0 | 20.9 |
